# In vitro modeling of nutritional and mitochondria-targeted therapies for osteosarcoma

**DOI:** 10.64898/2026.02.19.706776

**Authors:** Min Peng, Kelsey Keith, Shrey Dalwadi, Vernon E. Anderson, Adam Resnick, Marni J. Falk

## Abstract

Osteosarcoma is the most common pediatric bone tumor yet has limited treatment options, especially for metastatic cases with a 20% adjusted 5-year survival rate. Current therapies are non-specific, involving primary tumor resection with DNA-damaging chemotherapies like methotrexate, doxorubicin, and cisplatin. Few effective treatment options exist for metastases. Targeting metabolism involving cancer’s reduced mitochondrial functionality remains underexplored in osteosarcoma. We investigated the therapeutic potential in human osteosarcoma primary and metastatic cell lines of metabolic modulating drugs including metformin, cycloheximide, mitochondrial ETC inhibitors (antimycin A, metformin), dichloroacetate, and imipridones (ONC201, ONC206) on mitochondrial function and cell viability, individually and combined under various nutrient conditions across our lines. Results confirmed osteosarcoma cells are more dependent on glucose than osteoblasts but also require mitochondrial function for survival, highlighting the therapeutic potential of metabolic pathways. Osteosarcoma cell viability was reduced when any metabolic drug treatment was combined with conditions forcing reliance on mitochondrial OXPHOS capacity. Combination metabolic therapies, particularly ONC201/ONC206/metformin in 143B cells, and to a lesser extent DCA and ONC201 with either ONC206 or antimycin A, showed enhanced cytotoxicity compared to single agents, with a good therapeutic index based on minimal toxicity to normal osteoblast cells. The degree of effectiveness varied across cell lines, underscoring the importance of personalized treatment strategies. RNA-Seq transcriptome analysis revealed that effective nutrient and metabolic drug treatments triggered widespread regulatory changes in osteosarcoma cells involving increased translation/splicing with decreased mitochondrial processes such as cholesterol biosynthesis. These results demonstrate the utility of developing combined metabolic and chemotherapeutic treatments for osteosarcoma.

## INTRODUCTION

Osteosarcoma is the most frequent pediatric bone malignant tumor, with a peak incidence during teenage years (1,2). For children and adolescents, the incidence of osteosarcoma is 4.8 cases per million per year followed by a 5-year overall survival rate of approximately 70% (3). Unfortunately, 10-15% of osteosarcoma patients exhibit metastasis, primarily to the lungs. The adjusted 5-year survival rate for metastatic or recurrent osteosarcoma is 20%, highlighting a discrepancy in prognosis for aggressive tumors(4). Current treatment protocols include induction and consolidation therapies in tandem with surgical resection of the tumor. Chemotherapies commonly administered are methotrexate, doxorubicin (adriamycin), and cisplatin (platinol), also commonly referred to as the MAP regimen(5). These drugs broadly work through interference with cellular genetic material by damaging DNA and/or hindering DNA synthesis (6–8). Specific treatment guidelines for metastatic osteosarcoma are limited, though, relying on careful resection of spread tumor mass and delivery of similar drugs in clinical trials (5). For example, the dependence of sarcomas on activation of the CDKN2A-CCND-CDK4/6-Rb pathway has opened opportunities for the use of CDK4/6 inhibitors in the clinic (9), such as Abemaciclib (AMC, Verzenio®) used in the treatment of advanced or metastatic breast cancers and under study in osteosarcoma (10), to prevent G1/S cell cycle progression by blocking CDK4 and/or CDK6 from binding to their regulatory partner, cyclin D1 (CCND) (11) the inability to effectively target aggressive osteosarcomas justifies the need for an alternative therapeutic strategy.

One growing area of interest is understanding how tumor metabolism and nutrient availability can be harnessed to optimize chemotherapy. In the early 20^th^ century, Otto Warburg observed that cancer cells produced abnormally high levels of lactate even in the presence of oxygen. This shift in bioenergetics towards glycolysis is now commonly described as the Warburg effect (12). While the Warburg effect may seem to imply mitochondrial function is neither active nor necessary in cancer cells, recent studies emphasize that aerobic respiration and overall mitochondrial function is imperative to tumor growth (13). Consequently, interventions that modulate mitochondrial function are an emerging class of chemotherapeutics. For example, metformin, a common drug for type II diabetes, has been extensively studied in clinical trials for its anti-cancer potential (14). Although its mechanisms are complex and debated, metformin functions in part via indirect inhibition of mitochondrial respiratory chain (RC) complex I, activating AMP-activated protein kinase (AMPK) to ultimately inhibit mTORC1 and thereby decrease cellular proliferation (15). Recent brain tumor efficacy in diffuse midline gliomas (DMGs) and glioblastoma clinical trials has led to ONC201 (dordaviprone/Modeyso) being approved by the FDA for treatment of diffuse midline glioma harboring an *H3 K27M* mutation (16,17), highlighting the prospect of imipridones as a novel class of mitochondria-targeted chemotherapy (18). In the context of tumor metabolism, imipridones serve as agonists to caseinolytic protease P (ClpP), a serine protease that forms a heterodimer (ClpXP) with its chaperone, ClpX in the mitochondrial matrix (19). ClpXP plays a vital role in activating the unfolded protein response in mitochondria (UPR^mt^) via degradation of various misfolded peptides to restore mitochondrial function even in harsh tumor microenvironments (20). ONC201 and ONC206 are specific imipridones that exploit this behavior in tumors with increased ClpP expression by further hyperactivating CLPP, resulting in excessive degradation of essential RC complexes and ultimately cell death selectively in tumors, with surprisingly few adverse effects in healthy cells (19). While the success of imipridones has been demonstrated in numerous cancer types including ONC201’s recent FDA approval for the treatment of diffuse non-midline H3 K27M-mutant glioma, its therapeutic potential in osteosarcoma remains uninvestigated (19). However, many clinical attributes suggest their potential for efficacy. For one, osteosarcoma generally occurs near growth plates of the knee and shoulder joints, indicating a highly proliferative, and thus metabolically active, cell state (4). Moreover, resistance to the routine chemotherapeutic cisplatin may be correlated with elevated levels of ClpP, resulting in decreased cisplatin drug uptake (21). Therefore, shortcomings in current treatments, especially for aggressive osteosarcoma, highlight the need for an alternative approach that accounts for the dynamic metabolism of tumors.

Cycloheximide (CHX) is widely recognized as a potent inhibitor of protein synthesis by interfering with the cytosolic translation in mammalian cells. Several mechanistic effects of CHX have been reported in cancer cells, including inhibition of several actions of butyrate on human colorectal cancer cells including inhibiting the induction of alkaline phosphatase by butyrate in HT29 colon cancer cells (22), HRT-18 rectal cancer cells (23) and LoVo colon cancer cells (24). CHX also promotes paraptosis, which is a newly identified form of programmed cell death, characterized by dilation of the endoplasmic reticulum or mitochondria due to altered proteostasis or redox and ion homeostasis, that has been shown to be induced by inhibition of cyclophilins in glioblastoma multiforme (25). Low concentration CHX induced cell cycle arrest at G1 and S phases and cAMP-dependent differentiation in C6 glioma (26). Surprisingly, CHX at low micromolar concentrations also showed therapeutic benefit in primary mitochondrial disease models by preserving cellular respiratory capacity, inducing mitochondrial translation and biogenesis, and rescuing viability of human podocytes from autophagy, while also ameliorating proteotoxic stress induced by a range of RC inhibitors via a uniquely selective reduction of cytosolic protein translation (27). CHX also protected neuronal cells from oxidative stress (28). That CHX has therapeutic potential in both cancer and primary mitochondrial disease highlights the unique mechanistic intersection of metabolism and cancer.

Mitochondrial functional changes have been implicated in cancer. For example, RC complex III upregulation has been reported in the late stages of human high grade serous ovarian cancers (29). Interestingly, the antibiotic antimycin A (AA) is a potent complex III inhibitor whose application in anticancer studies is rare although recently ovarian (29) and liver (30) cancer cells with high complex III subunit expression have shown high sensitivity to AA (29,30). Further, targeting mitochondrial complex III and OXPHOS has been shown to eradicate cancer stem cells (31). In addition, pyruvate dehydrogenase complex (PDHc) is a key mitochondrial enzyme that has been recognized as a key metabolic node in cancer with upregulated activity in osteosarcoma (32). Dichloroacetate (DCA), which activates PDHc via inhibiting the PDH kinase that phosphorylates and thereby reduces PDHc activity, has long been suggested to have potential anticancer effects in preclinical or clinical trials (33–35).

Here, osteosarcoma cell viability was evaluated when grown in media with different nutrient compositions involving low or medium glucose concentrations, or galactose. The preclinical efficacy on cell viability of imipridones (ONC201, ONC206), metformin, CHX, and AA, were evaluated in seven human osteosarcoma cell lines (6 primary tumor derived lines: 143B, HOS, MG-63, SAOS-2, SJSA-1, U-2 OS; a single pulmonary metastasis derived line: 15454-307) in different nutrient concentrations and treatment durations. Imipridone effects were further evaluated by ClpX/P immunoblot expression analyses, and RNA-Seq transcriptome profiling was performed to dissect the cellular mechanisms of nutrient and metabolic drug treatments in human osteosarcoma.

## MATERIALS and METHODS

### Subject

Sex as a biological variable was not considered as a variable in this paper. However, all cell lines used were from donors with a mix of sexes. The ages of the original cell line donors were primarily in their teens, which is the most common age for development pediatric osteosarcoma.

### Bias reduction

As no human subjects were involved in this study, no bias reduction strategies such as blinding or randomization were utilized and there were no inclusion or exclusion criteria.

### Cell Line Description and Culture Conditions

Authenticated human osteosarcoma cell lines were obtained: 143B (ATCC Cat# CRL-8303, RRID:CVCL_2270), HOS (ATCC Cat# CRL-1543, RRID:CVCL_0312), MG-63 (ATCC Cat# CRL-1427, RRID:CVCL_0426), Saos-2 (ATCC Cat# HTB-85, RRID:CVCL_0548), SJSA-1 (ATCC Cat# CRL-2098, RRID:CVCL_1697), U-2 OS (ATCC Cat# HTB-96, RRID:CVCL_0042) and osteoblast cell lines hFOB 1.19 ((ATCC Cat# CRL-3602, RRID:CVCL_3708), hFOB) and NHOst (Lonza Cat #: CC-2538). Osteosarcoma cell line 15454-307 was established from a pulmonary metastasis from an osteosarcoma patient isolated at Children’s Hospital of Philadelphia (IRB # 8-015454, AR PI). Q2148 is a healthy control fibroblast cell line established from the same osteosarcoma patient. Q1875p1 is a transmitochondrial cybrid line of 143B cells with mtDNA genome from a healthy individuals’ fibroblast cell line. Both fibroblast lines were consented as part of the ongoing CHOP Mitochondrial Medicine Frontier Program (MMFP) study (IRB #18-015454). Cells were cultured at 37℃ with 5% CO_2_ either in DMEM-10% FBS, 0.2% uridine (15 µM) with 5.6 mM glucose, 25 mM glucose, or 10 mM galactose; or DMEM / F12 −10% FBS 17 mM glucose. Upon reaching 80-90% cell confluency, cells were passaged and counted using a TC20 Automated Cell Counter (Bio-Rad, Hercules CA).

### Drugs Studied

Compounds were obtained as follows: CHX (Sigma-Aldrich, St. Louis MO, Cat# C19865), metformin (MP Biomedicals, Irvine CA, Cat# 151691), imipridones ONC201 and ONC206 (Selleckchem, Cat#s S796301 and S6853,respectively; are no longer available from this source but are now available from Cayman Chemical), dichloroacetate (DCA, Sigma, CAT# 347795), and AA (Sigma CAT# A8674). Stock solutions of all drugs were prepared in DMSO.

### Cell Survival Assay

After reaching 80-90% confluency, cells were seeded at 10,000, 5,000 or 2,500 cells/well of 96-well plates (Corning, Corning NY, Cat# 3585 with 100 µL/well of DMEM-10% FBS-25 mM glucose or DMEM/F12-10% FBS 12.5 mM glucose and incubated at 37℃ . After either 5 h or overnight, the media was exchanged with 100 µL/well or 200 µL/well of fresh media (25 mM, 12.5 mM, 5.6 mM glucose, or 10 mM galactose) containing one of the identified drugs or DMSO as buffer only control. The plate was then incubated for one of the tested treatment durations (24, 48, or 72 h, or 6 days). At the end of the treatment, an MTT Cell Proliferation Assay Kit (Abcam, Cambridge UK, Cat# ab211091) was used to quantify cytotoxicity. The media in each well was exchanged for 100 µL/well of DMEM with MTT reagent, and the plates were incubated at 37°C for 4 h. 150 µL/well of stopping solution was then added, and plates were placed on an orbital shaker for 15 minutes under foil at room temperature. Absorbance was measured at 590 nm in a microplate reader and normalized to the control group. For intergroup comparisons, hFOB was used as the control for tumor lines, untreated buffer-only exposed tumor lines were used as controls for treatments in the same lines, and 5.6 mM glucose was used as the control for alternative nutrient conditions.

### Mitochondrial Physiology Analysis by High Content Screening (HCS) Assay

The CellInsight® CX5 HCS Platform (ThermoFisher, Waltham MA) was used to analyze both the mitochondrial membrane potential and mitochondrial content. 2,500 cells per well were seeded into a 96-well plate with DMEM-10% FBS 25 mM or 5.6 mM glucose overnight, after which media was replaced with 100 µL per well of the same medium containing diamidino-2-phenylindole (DAPI) as NucBlue® Fixed Cell ReadyProbes® Reagent (ThermoFisher, Cat# R37606) plus mitochondrial membrane potential probe, tetramethylrhodamine methyl ester (TMRM) (ThermoFisher, Cat# #T668), or mitochondrial content probe (MitoTracker Green, ThermoFisher Cat# M46750) at 37℃ for 20 minutes. The staining solution was aspirated, cells were rinsed with phosphate buffered saline (PBS), and images were acquired on the CellInsight®CX5 HCS Platform with a 10× objective in two different channels. Cell nuclei were identified and counted using the DAPI fluorescence 350/460 nm (excitation/emission) wavelengths (36). Fluorescence from cell debris and other particles was excluded using a size filter tool. The pixel intensity of the “spots” with the region of Interest (ROI) based on nuclei and either TMRM (548/574 nm) or MitoTracker Green (490/516 nm) fluorescence was identified, counted, and measured by ThermoFisher Scientific® HCS Studio® Cell Analysis Software.

### Western Immunoblot Analyses

143B and hFOB cells were seeded onto 100 mm Petri dishes overnight to reach 80-90% confluence. Drug solutions (10 mL) were created with DMEM (1 g/L) and various ONC201 and ONC206 concentrations (DMSO control). At desired timepoints, plates were placed on ice for cell lysis. Cells were washed twice with cold PBS, scraped into a 15 mL tube, and centrifuged at 500×*g* for 5 minutes at 6°C. Supernatant was aspirated and lysis buffer (radioimmunoprecipitation assay buffer (RIPA), Sigma, Cat# #20-188) with phenylmethanesulfonyl fluoride (PMSF, 1% v/v Sigma #329-98-6) and Protease Inhibitor Cocktail (Sigma-Aldrich, CAT# p8340) 1% v/v was added to the cell pellet, thoroughly mixed, and transferred to a microcentrifuge tube. The solution was kept on ice for 30 minutes with regular mixing. Lysis solution was centrifuged at 20,000×*g* for 10 minutes at 4°C. SDS-PAGE loading buffer (5×) was added at 1:5 ratio to the supernatant of the lysis solution. After vortexing, proteins were denatured in a heat block (97°C) for 4 minutes and then stored in a - 20°C freezer. Equal volumes of protein solutions were added to the wells of a mini-PROTEAN TGX Precast Gel (4-15%, Bio-Rad Laboratories) along with a molecular weight marker lane. Running buffer was filled to the top of the cassette in the chamber and the gel was electrophoresed at 70 V until the sample entered the gel. Then, the voltage was increased to 95 V for about 1 h (or until the dye front was eluted). Afterwards, the gel was removed from the cassette and placed inside a transfer stack with nitrocellulose paper covered in transfer buffer. This stack was moved to another chamber, pre-cooled with ice and transfer buffer. The transfer process was run for 1 h at 350 mA. The nitrocellulose membrane was then washed and incubated in blocking buffer for 2 h. Primary antibodies (Cell Signaling Technology) against ClpP (Cell Signaling Technology Cat# 14181, RRID:AB_2798414), ClpX, citrate synthase (CS), and glyceraldehyde-3-phosphate-dehydrogenase (GAPDH) were incubated overnight at 4°C at the recommended dilution (1:1000). After washing 3× with Tris buffered saline (TBS), the membrane was incubated with an appropriate secondary antibody (1:10,000) for 1 h at room temperature and washed 4× with TBS with 0.1% Tween and once with TBS. Immunoblot images were digitized utilizing a fluorescence scanner at appropriate wavelengths, intensities integrated in ImageJ (RRID:SCR_003070) and then normalized to either CS or GAPDH as loading control and then to control samples, normalized intensity = (I_ClpP/X(drug)_/I_CS/GAPDH(drug)_) / (I_ClpP/X(cntrl)_/I_CS/GAPDH(cntrl)_).

### RNA-Seq Transcriptome Profiling

Cells in 100 mm dishes (about 80% confluency) were replaced with DMEM-10% FBS 5.6 mM glucose, or 10 mM galactose, or 10 mM galactose plus drug (0.5 µM CHX, 1.3 µM ONC201, 0.12 µM ONC206, or 2.5 mM metformin). After culturing for 24 h, cells were washed with cold PBS twice and pelleted in a 1.5 mL tube. Cells (about 1 million) were lysed with 1 mL TRIzol® reagent (Invitrogen Cat# 15596026) and homogenized by pipetting up and down several times. RNA was purified with RNeasy Kits (Qiagen, Hilden Germany, Cat# 74004). RNA-Seq data were processed with the nf-core RNA-sesq pipeline (37).

Briefly, the nextflow pipeline processes the data by aligning (38,39) . Additional details are available in the (17,40–42) for differential expression, with (43) for pathway analysis using the Reactome (44) to calculate normalized enrichment scores (NES) and plots were created with custom scripts using the tidyverse, especially ggplot2 (45,46) For looking at overlaps in gene sets of interest after ONC201 and ONC206 treatment, over-representation analysis (ORA) with WebGestaltR (47) was used.

### Statistical Analysis Methods

All non-sequence data were plotted and analyzed using R (RRID:SCR_001905). Unless otherwise noted, each experiment was performed with three biological replicates. Plots were made using ggplot2 (RRID:SCR_014601). Comparisons between groups were analyzed using ANOVA with a post-hoc Tukey test and blocking for replicates. An adjusted probability (P) value of less than 0.05 was considered statistically significant. All code for RNA-seq and statistical analysis available from authors upon request.

## RESULTS

### Osteosarcoma cell proliferation, mitochondrial function, and transcriptome adaptations are nutrient dependent

Nutrient effects on osteosarcoma cell line 143B viability relative to hFOB osteoblast control cells were studied following 24-h culture in low glucose (5.6 mM) or high glucose (25 mM) media. Clear glucose-dependence was seen for 143B cell viability, which exhibited a 2-fold increase in low glucose when normalized to hFOB cells in three biological replicate experiments (P < 0.001, **Figure 1A**) which increases to almost 3-fold enhanced proliferation in high glucose. As predicted by the Warburg effect, these results are consistent with overreliance of osteosarcoma cells on anaerobic glycolysis. Therefore, we next sought to determine if dysfunctional mitochondria in osteosarcoma cells underlie the need for increased glycolytic flux, where mitochondrial membrane potential was used as an indicator of integrated mitochondrial RC function. Interestingly, mitochondrial membrane potential of 143B osteosarcoma cells was elevated in high glucose conditions and reduced in low glucose conditions relative to hFOB osteoblast cells in three biological replicate experiments (P < 0.001, **Figure 1B**), despite the presence of similar mitochondrial mass in both osteoblast and osteosarcoma cells (**Figure 1C**). Culturing cells in galactose media without glucose present is a common experimental strategy to force cells to survive off their mitochondrial function (48), as galactose cannot be as effectively used by glycolysis to generate cellular ATP. As expected, 143B osteosarcoma cells grown in 10 mM galactose media for 72 h had significantly decreased viability relative to their growth in glucose media, again suggesting osteosarcoma cells have dysfunctional mitochondria since cells with healthy mitochondria have no change in viability when grow**n** in galactose media (**Figure 1D**). Viability in galactose media relative to low glucose media for 3 days was similarly reduced in 4 additional osteosarcoma primary cell lines (MG-63, Saos-2, SJSA-1, U-2 OS), and a pulmonary metastatic osteosarcoma patient-derived cell line (15454–307) in three biological replicate experiments (P < 0.0001, **Figure 1E**). Interestingly, the HOS osteosarcoma cell line did not have reduced viability in galactose media. Overall, these in vitro data confirm the predominant finding of glucose dependence, with reduced mitochondrial RC function, in primary and metastatic osteosarcoma cells.

**Fig 1.**
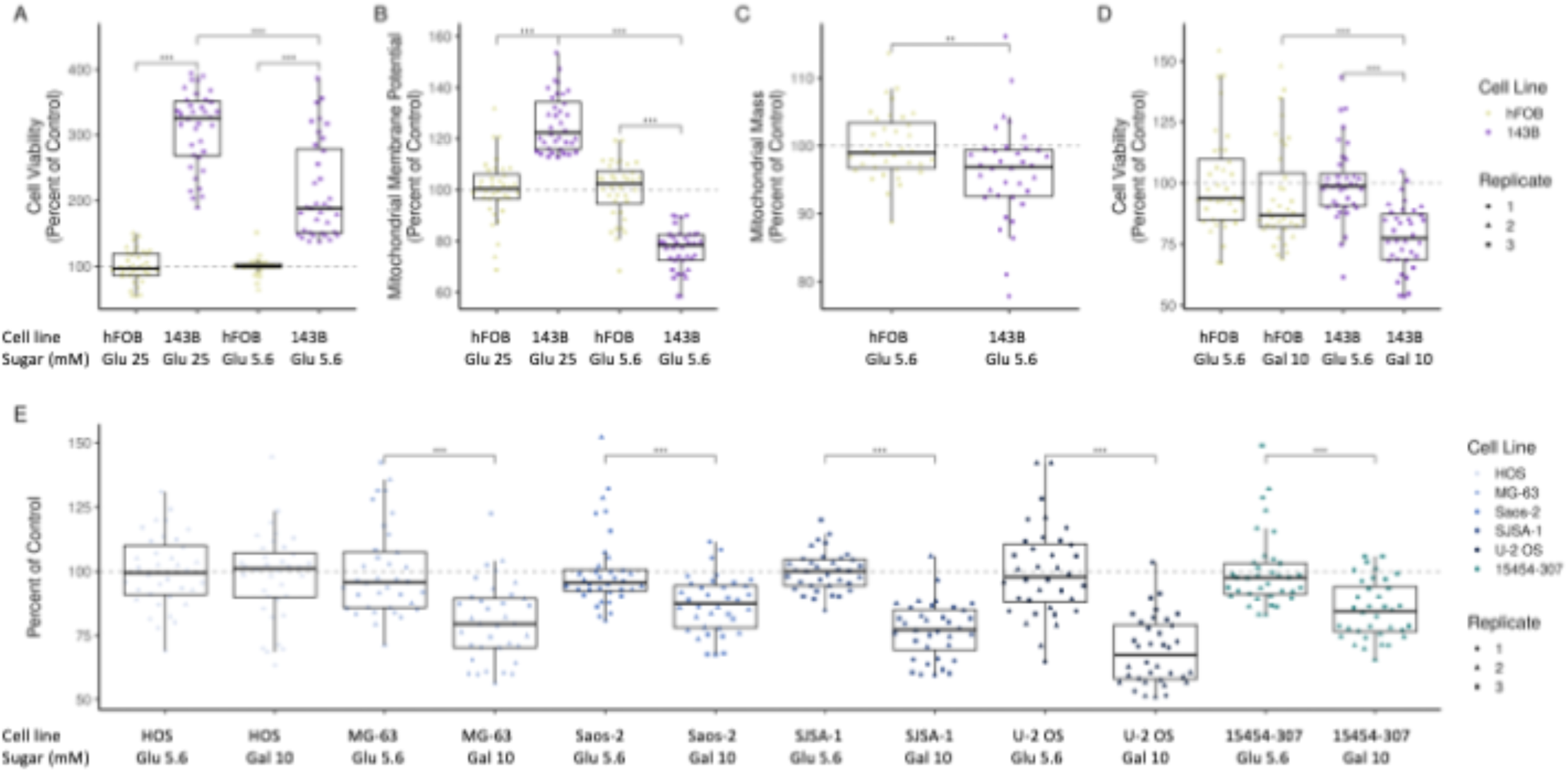
Cell viability and mitochondrial function of osteosarcoma cells rely more on glucose compared to osteoblast cells. **(A)** 143B cells (osteosarcoma) show greater proliferation than hFOB cells (osteoblast control cells) in both high (25 mM) and low (5.6 mM) glucose media in three replicate experiments. **(B)** Mitochondrial membrane potential of 143B cells was significantly decreased when glucose media was changed from 25 mM to 5.6 mM comparing with hFOB cells in three biological replicate experiments.. **(C)** Mitochondrial membrane potential of the osteosarcoma cells was surprisingly greater in high glucose but lower in low glucose media. **(D)** Cell viability of 143B cells normalized to hFOB cells in 5.6 mM glucose or 10 mM galactose medium. Only the 143B osteosarcoma cells had significantly reduced viability in galactose media compared to both hFOB cells in galactose and 143B cells in glucose. **(E)** Cell viability of 6 osteosarcoma cell lines (including 5 primary tumor derived lines: HOS, MG-63, Saos-2, SJSA-1, U2 OS, and one pulmonary metastasis derived line: 15454-307) grown in 5.6 mM glucose or 10 mM galactose medium. Significant reductions of cell viability to variable degrees were seen in galactose conditions for all cell lines except HOS. Three biological replicate experiments per condition. All boxplots show the median of the data, the boxes mark the limits of the second and third quartiles and the whiskers the smallest and largest values no more than 1.5 times +/- the interquartile range. Points outside that range are considered “outlying” and are plotted individually. Differences between the groups were tested with ANOVA with a post-hoc Tukey test while controlling for biological replicates.

Mechanistic evaluation of reduced osteosarcoma proliferation in galactose relative to glucose media was performed using RNA-Seq transcriptome profile on the same 7 osteosarcoma cell lines (primary osteosarcomas 143B, HOS, MG-63, Saos-2, SJSA-1, U-2 OS and pulmonary metastasis-derived patient osteosarcoma cell line 15454-307). Each cell line showed similar proportions of ∼2% differentially expressed genes in response to the different bioenergetic energy sources (**Figure 2A,C“,E,G,I,K,M**). While the magnitude of changes was similar with some universal responses seen, each osteosarcoma cell line also displayed unique responses upon growth in galactose (**Figure S1A**). Universal changes mainly corresponded to upregulated pathways involved in RNA transcript handling, such as rRNA processing, mRNA editing, and RNA polymerase transcription; upregulated interferon response was also seen in two osteosarcoma cell lines grown in galactose media. Mitochondrial metabolism pathways including cholesterol biosynthesis, and fatty acid synthesis, were all downregulated in galactose media. Interestingly, galactose incubation caused the metastatic osteosarcoma cell line 1554-307 to upregulate cell cycle pathways, signaling cell cycle arrest (**Figure 2N**). Despite some common shifts, each osteosarcoma cell line also had unique responses upon a switch to galactose from glucose media. For example, SJSA1 upregulated CYP drug metabolism pathways; Saos-2 upregulated TP53 pathways; and 15454-307, in addition to the cell cycle arrest, demonstrated transcriptional responses consistent with defective DNA repair. Surprisingly, some cell lines responded by regulating the same pathways in completely opposite directions, such as FGFR fibroblast receptor pathways that were upregulated in HOS and downregulated in U-2 OS osteosarcoma cells. Overall, transcriptome profiling reflected the known genetic heterogeneity of osteosarcomas from different individuals.

**Fig 2.**
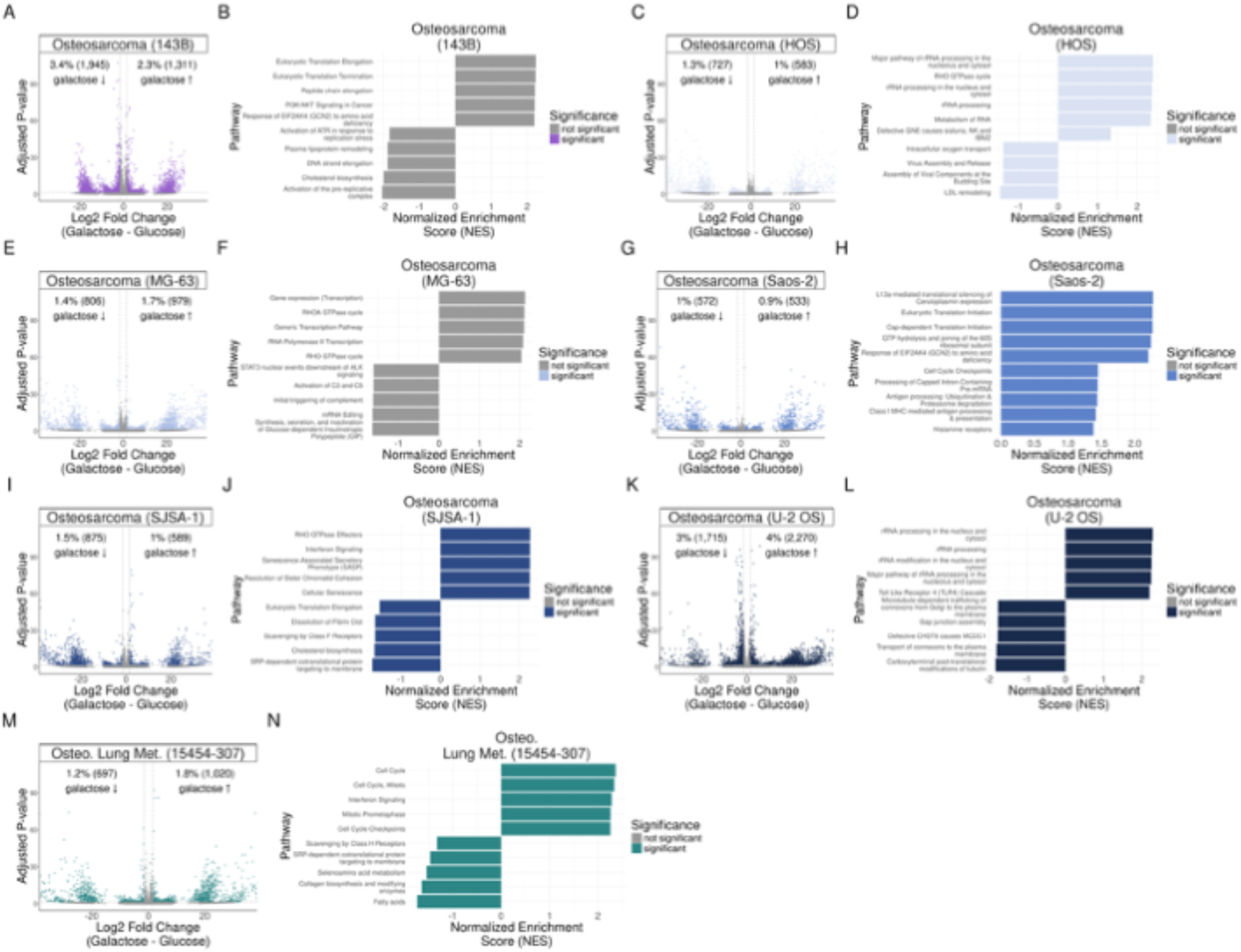
Osteosarcoma cell line RNA-Seq transcriptome profiling in 7 osteosarcoma cell lines. Differential expression, tested using DESeq2, and pathway analysis, tested using fgsea, is shown for 143B (A, B), HOS (C, D), MG-63 (E, F), Saos-2 (G, H), SJSA1 (I, J), U-2 OS (K, L) and osteosarcoma lung metastasis 15454-307 (M, N) osteosarcoma cell lines. Volcano plots display differential expression results with log2 fold change on the x-axis and negative log10 adjusted p-value on the y-axis (A, C, E, G, I, K, M). Bar plots detail the top five most up- and down-regulated pathways (B, D, F, H, J, L, N). Both significantly different points in the volcano plot and significant pathways in the bar plots are colored by cell line; non-statistically significant points or bars are shown in gray. Legends are grouped, with the legend applying to both the volcano plot and the bar plot of the same color.

### CHX, metformin, ONC201, or ONC206 single or combined metabolism-targeted therapies inhibited osteosarcoma 143B cell growth and/or viability in glucose media

*In vitro* metabolism-targeted drug testing in low glucose conditions of metformin, ONC201, or ONC206 treatment for 72 h was conducted utilizing an MTT assay to quantify cell viability. Metformin at 2.5 mM and 5 mM concentrations significantly decreased cell viability by 67% and 89%, respectively in 143B osteosarcoma cells, with no effect of 2.5 mM metformin and the only significant, but mild, inhibition of cell viability occurred with 5 mM metformin in hFOB osteoblasts (P < 0.001, **Figure 3A**). The imipridones induced decreased viability of 143B cells grown in glucose media by 52% with 1.3 µM ONC201 (P < 0.001) and by 57% with 0.06 µM ONC206 (P < 0.001) relative to untreated 143B cells in three biological replicate experiments, with a statistically significant, but small effect on hFOB cell viability with ONC201 only (**Figure 3B,C**). To evaluate for a synergistic effect of metabolic-targeted treatments, combinatorial drug regimen effects on osteosarcoma cell viability were explored. Combining 2 mM metformin and either 0.26 µM ONC201 or 0.06 µM ONC206 (**Figure 3D**) in three biological replicate experiments resulted in selective cell cytotoxicity for 143B osteosarcoma cells to a respective mean of 16% or 33% viability relative to untreated 143B cells in 5 mM glucose media for 3 days, with no effect on hFOB osteoblast cells. Interestingly, 143B cells treated simultaneously with both imipridones (0.65 µM ONC201 and 0.06 µM ONC206) consistently and significantly decreased 143B survival but to variable degrees in 3 biological replicate experiments, suggesting possible synergy may exist when combining these two imipridone molecules in osteosarcoma, with a more modest effect in osteoblasts (P < 0.05, **Figure 3D“)**. However, 3-way treatment of 2.5 mM metformin plus low doses of both imipridones (0.65 µM ONC201 and 0.06 µM ONC206) reduced 143B cell viability to ∼10% relative to untreated cells (P < 0.001), with a partial inhibitory effect to only ∼50% viability for the hFOB cells (P < 0.001) **(“Figure 3D“).** CHX effects at low concentrations (0.1, 0.25 or 0.5 µM) were studied on viability **(“Figure 3E“)** and mitochondrial membrane potential **(“Figure 3F“)** of 143B and hFOB cells grown in low glucose medium. CHX significantly but partially reduced both cell viability (P < 0.001) (**Figure 3E**) and mitochondrial membrane potential (P < 0.001) (**Figure 3F“)** in 143B cells after a 72-hour CHX treatment in a dose-dependent manner in three biological replicate experiments, with no significant effect on hFOB cell viability.

**Fig 3.**
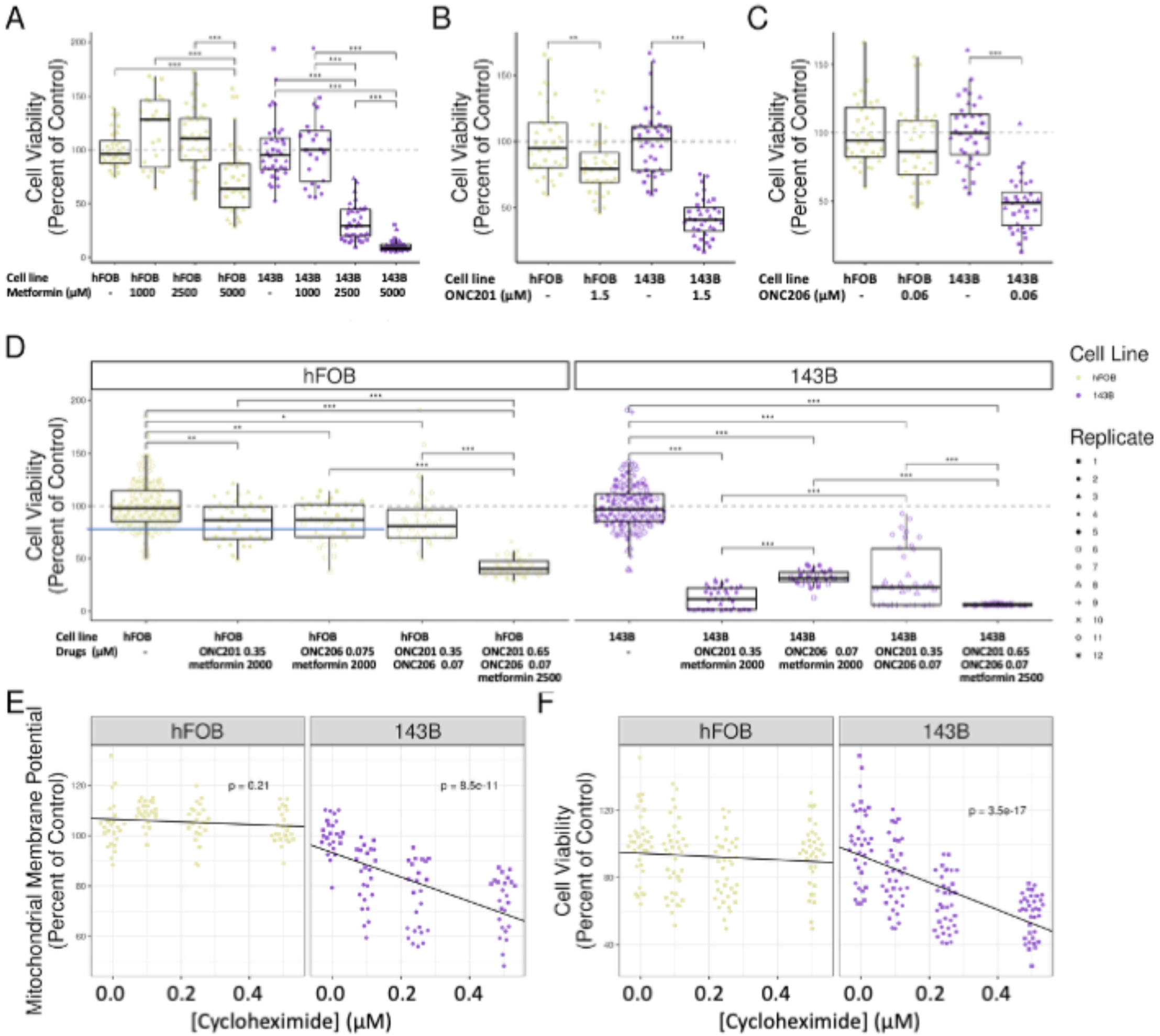
Metformin, cycloheximide, ONC201, ONC206, or their combinations suppressed osteosarcoma 143B cells growth but not osteoblast hFOB cells in low glucose (5.6 mM) medium. 2500 cells per well of 143B cells or hFOB cells were seeded in 96 well in 5.6 mM glucose DMEM-10% FBS medium, and 5 h later, treated with **(A)** metformin, **(B)** ONC201 or **(C)** ONC206, or **(D)** a combination of metformin, ONC201, and ONC206. MTT cell viability assays were run after culturing for 72 h and the differences in cell viability were tested using ANOVA with a post-hoc Tukey test, n = 3 biological repeats. **(E)** Low dose of cycloheximide (CHX) reduced 143B cell mitochondrial membrane potential and **(F)** suppressed 143B cells growth vs hFOB cells. 2500 cells per well of 143B or hFOB cells were seeded in 96 well in 5.6 mM glucose DMEM-10% FBS medium and 5 h later CHX was added. The mitochondrial mass assays were run after 24 h culturing on a CellInsight CX5 HCS platform and MTT cell viability assays were run after culturing for 72 h with n = 2 or 3 biological replicates. Results were tested with a linear model for CHX concentration vs cell viability or mitochondrial membrane potential respectively **(D, F)** controlling for batch. Points in the linear models are jittered to represent the underlying distribution of the data and increase its visibility; all models were fitted on original data.

To evaluate the broader impact of these metabolic treatments across diverse osteosarcoma cell lines, optimal treatment conditions identified in 143B osteosarcoma and hFOB osteoblast experiments were subsequently screened in 1 or 2 biological replicate experiments in a wider array of osteosarcoma (143B, Q1875p1, 15454-307, HOS, MG-63, Saos-2, SJSA-1, and U-2 OS) and control cell lines (osteoblasts hFOB and NHOst, and fibroblast Q2418). Specifically, cells were treated with 5 µM CHX for 6 days (**Figure 4A-D“, Figure S2A-D**) or 2.5 mM metformin, 1.3 µM ONC201, or 0.12 µM ONC206 for 72 h in either low glucose (5.6 mM, **Figure 4 E-I**, **Figure S2E-H**) or moderate glucose (17 mM, **Figure 4 J-N**, **Figure S2 I-N**) and DMEM-10% FBS conditions. CHX treatment for 6 days significantly decreased cell viability in all cell lines (p < 0.001), both in healthy osteoblasts hFOB (**Figure 4A**), and in 7 osteosarcoma lines including 143B (**Figure 4B**), MG-63 (**Figure 4C**), 15454-307 (**Figure 4D**), HOS (**Figure S2A**), Saos-2 (**Figure S2B**), SJSA1 (**Figure S2C**), and U-2 OS (**Figure S2D**). Interestingly, 2.5 mM metformin treatment did not significantly impact most of the osteosarcoma cell lines’ viability in low glucose conditions except for Q1875p1 (cybrid of 143B nucleus with normal fibroblast mitochondria, **Figure 4E**, p < 0.001) and U-2 OS cells (**Figure 4F**, p < 0.001). However, the imipridones ONC201 and ONC206 significantly, albeit modestly, reduced viability in <u>low glucose</u> media of HOS by 35%, p < 0.001, and 41%, p < 0.001, respectively (**Figure 4G“)**; Saos-2 by 50%, p < 0.001, and 49%, p<0.001, respectively (**Figure 4H“);** SJSA1 by 43%, p <0.01, and 47%, p< 0.01 respectively **(“Figure 4I“)**, tumor cancer cell lines. By comparison, no toxic effect of the imipridones were seen in either of two osteoblast cell lines (hFOB **Figure 4D**, NHost **Figure S2E**) or healthy human fibroblasts (Q2148, **Figure S2F**) studied. Notably, the greater inhibitory effect on cell viability of imipridones relative to metformin *in all cases* highlights the potential value of a mitochondria-targeted therapy. Interestingly, the imipridones induced no reduction of viability in either control or osteosarcoma cells when grown in <u>moderate glucose</u> (17 mM) conditions **(“Figure 4 J-M“),** with the exception of a significant modest (25%, p<0.01) reduction in viability by ONC201 in HOS and Saos-2 cell lines (**Figure S2L**, **Figure 4N**). These data suggest that ample glucose conditions protect osteosarcoma viability from mitochondria-targeted therapeutics.

**Fig 4.**
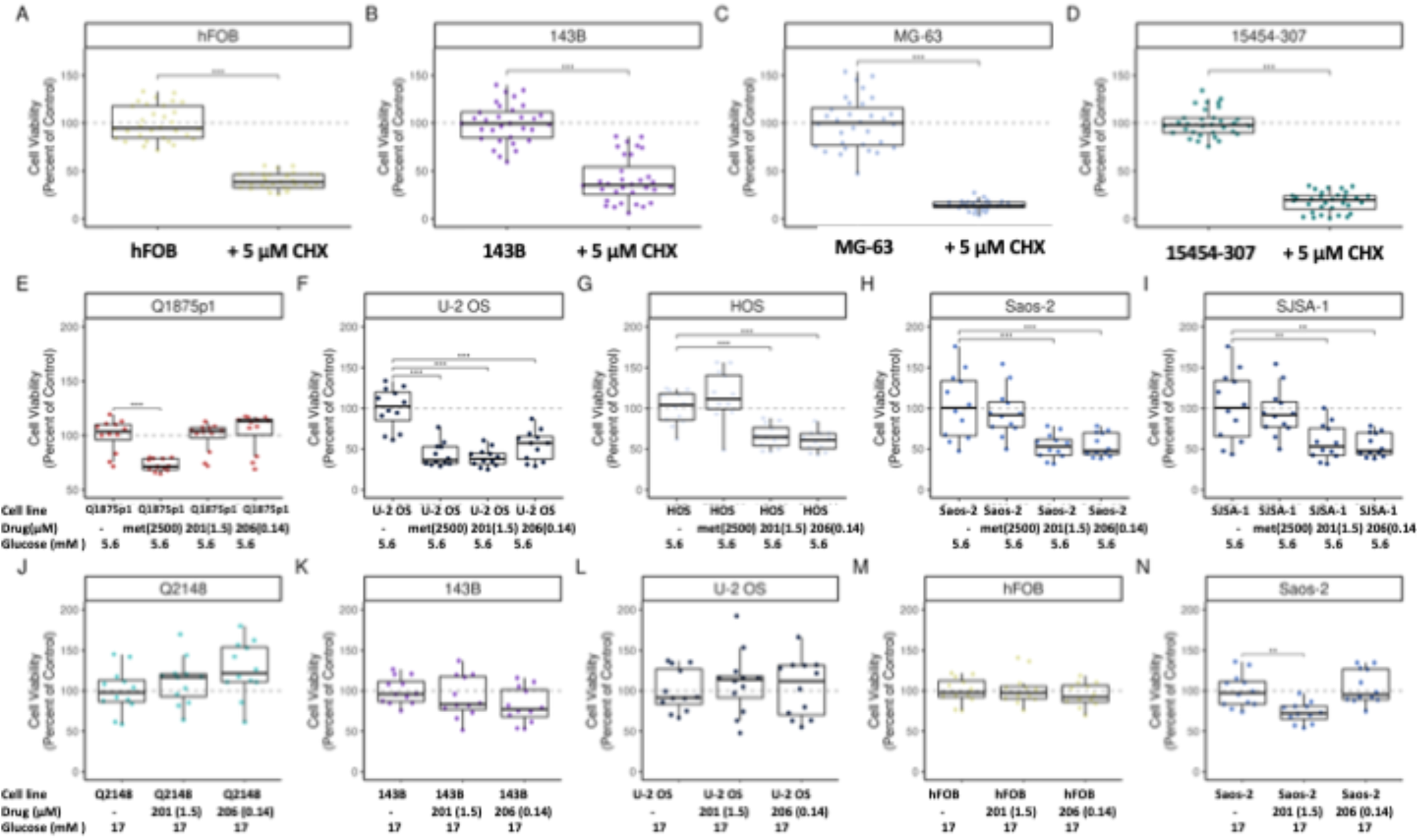
Efficacy of single agent treatment with cycloheximide, metformin, ONC201 and ONC206 on healthy cells, (hFOB and NHOst osteoblasts and normal Q2148 fibroblasts vs osteosarcoma cells (143B, MG-63, and 15454-307). Cell viability of **(A)** healthy osteoblasts hFOB, **(B)** primary osteosarcomas 143B and **(C)** MG-63, and **(D)** lung metastasis 15454-307 treated with 5 µM cycloheximide. Cell viability of **(E)** healthy osteoblasts NHOst, **(F)** healthy fibroblasts Q2148, and osteosarcomas **(G)** MG-63 or **(H)** 15454-307 treated with metformin, ONC201, or ONC206 in 5.6 mM glucose DMEM-10% FBS medium for 72 h. Cell viability of the same control cells and osteosarcoma cells, **(I)** hFOB, **(J)** NHOst, **(K)** Q2148, **(L)** 143B, **(M)** MG-63, **(N)** 15454-307 treated with 1.5 µM ONC201, or 0.15 µM ONC206 in 17 mM glucose DMEM-10%FBS medium for 72 h. Cell viability was tested using ANOVA with a post-hoc Tukey test for all comparisons. Box plots identify the median second and third quartiles and the whiskers extend to the smallest and largest values no more than 1.5 times +/- the interquartile range. Points outside that range are considered “outlying” and are plotted individually.

To investigate the observed differences in viability effects of metabolic treatments in primary and metastatic osteosarcoma cell lines, RNA-Seq transcriptome profiling was performed on the 143B primary osteosarcoma cell line and 15454-307 metastatic osteosarcoma cell line treated with reduced dosages 2.5 mM metformin, 1.3 µM ONC201, 0.12 µM ONC206, or 0.5 µM CHX grown in 10 mM galactose for 48 h, instead of 72 h, in order to limit toxicity to obtain sufficient live cells for RNA-Seq (**Figure 5**). In both cell lines, ONC201 similarly affected translation initiation, selenoamino acid metabolism, and nonsense-mediated decay in both 143B primary and 15454-307 metastatic cell lines. However, ONC201 uniquely upregulated DNA damage reversal and DNA replication in the metastatic line while NTRK signaling, part of the MAPK pathway was upregulated in both, and replication and DNA repair, specifically base excision repair, were uniquely downregulated in the primary osteosarcoma. ONC206 changed the expression of fewer genes in both the primary (5% ONC201 vs 2.9% ONC206) and metastatic (4.3% ONC201 vs 2.8% ONC206) cell lines, suggesting that ONC206 may have more targeted effects as it affected fewer genes. Even though they are analogues, for those genes that were differentially expressed, only ∼1/3 were commonly changed between the cell lines after ONC201 treatment **(Figure S1B)**. Several similar pathways were affected in the primary cell line with ONC201 and ONC206 treatment, relating to mRNA translation initiation and processing and upregulation of NTRK signaling. Conversely, there were no significantly different pathways with ONC206 in the metastatic cell line, demonstrating again that the analogues do not have equivalent effects. Interestingly, CHX treatment upregulated developmental pathways involving HOX genes and both WNT signaling as well as receptor tyrosine kinase signaling, all more strongly in the metastatic vs the primary cell line. Conversely, CHX upregulated the cell cycle in both cell lines, but to a greater degree in primary vs metastatic osteosarcoma cells as well as upregulating NTRK signaling. Uniquely in 143B, CHX upregulated mTORC1 as well as cell surface signaling and communication. Uniquely in 15454-307, CHX upregulated AMPK signaling and downregulated PALB2 function in homologous recombination repair (**Figure 5L“)**. Finally, metformin changed a similar number of genes in both the primary 143B (10.9%) and metastatic 15454-307 (11.1%) cell lines (**Figure 5M“, O**). In 143B cells, it upregulated RNA translation and processing pathways, similar to treatment with either imipridone.

**Fig 5.**
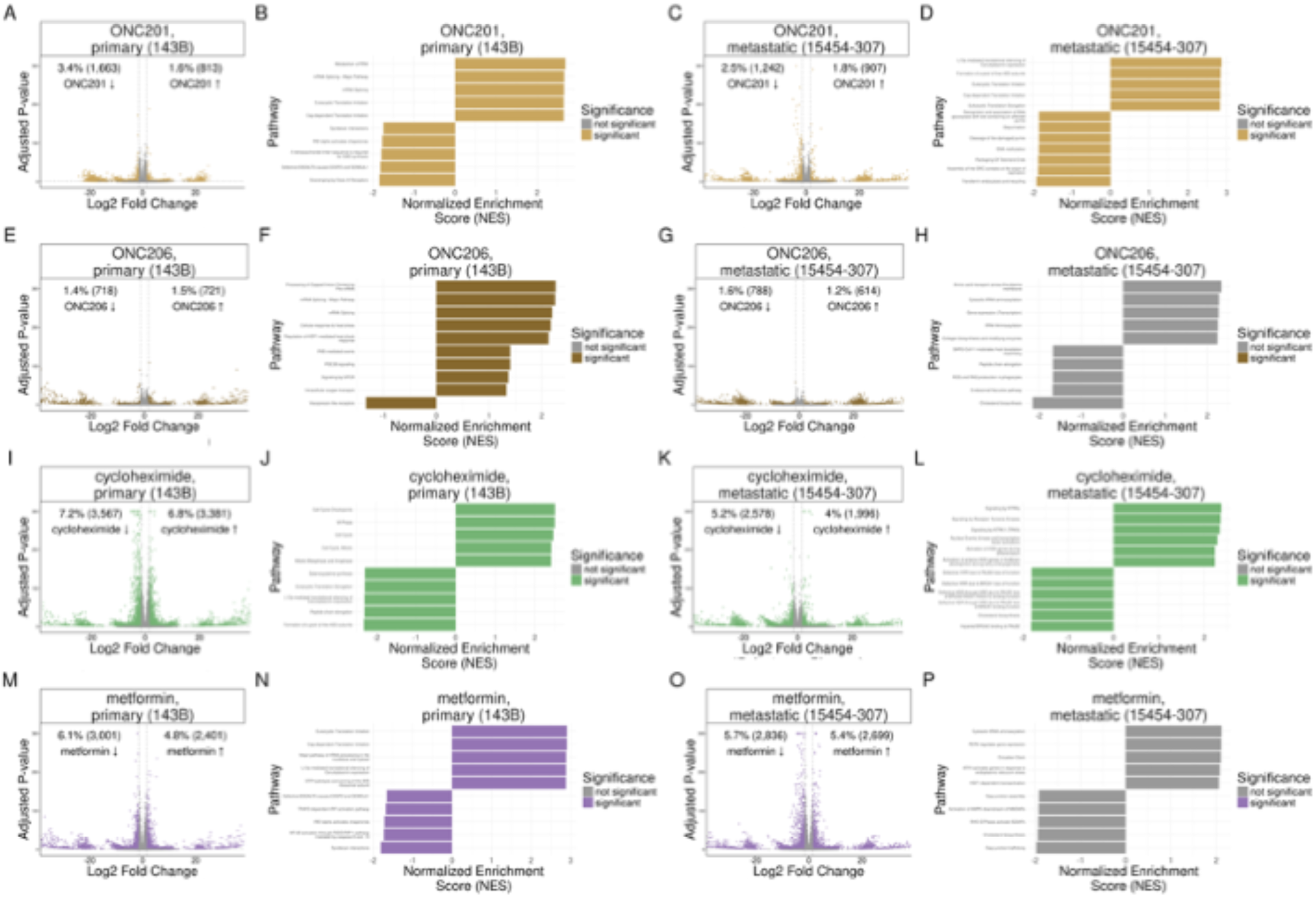
Osteosarcoma cell line RNA-Seq transcriptome profiling of differential expression and pathway analysis in 143B (A-H), 15454-307 (I-P) osteosarcoma, cell lines treated with 1.3 µM ONC201 (A, B, I, J), 0.12 µM ONC206 (C, D, K, L), 0.5 µM **cycloheximide (E, F, M, N) or metformin (G, H, O, P).** Volcano plots show differential expression results, tested using DESeq2, with log2 fold change on the x-axis and negative log10 adjusted p-value on the y-axis (A, C, E, G, I, K, M, O). Bar plots give the top five most up- and down-regulated pathways, (B, D, F, H, J, L, N, P) tested using fgsea. Both significantly different points in the volcano plot and significant pathways in the bar plot are colored by treatment; non-statistically significant points or bars are shown in gray. Legends are grouped, with the legend applying to both the volcano plot and the bar plot of the same color.

Downregulated pathways were different however with TRAF3-depent IRF activation and NF-kB inhibited. Like ONC206 treatment, metformin treatment did not significantly change any pathways in 15454-307. As upregulation of NTRK signaling was common across cell lines and treatments, this may represent a common escape mechanism from chemotherapy treatment, as overexpression of any of NTRK1-3 is known to drive proliferation in a wide variety of both pediatric and adult cancers, making NTRK inhibitors an interesting approach for future work (49). Additionally, this diversity of response to treatments suggested that mitochondria-targeting combination therapy will be more effective than single-agent therapy as drugs will affect different pathways, leading to synthetic lethality.

### Prolonged and combinatorial metabolic-targeted treatments selectively induced greater impairment of osteosarcoma viability

To evaluate the comparative efficacy of mitochondrial metabolism-targeted drugs for prolonged duration, in vitro viability studies over 6 days of all 7 osteosarcoma cell lines relative to hFOB osteoblasts were performed in high glucose (17 mM) media individually with AA, DCA, metformin, ONC201, or ONC206, or in select combinations. 25 nM AA had no significant effect on hFOB cell viability (**Figure 6A**) while reducing cell viability by 37% in 143B (**Figure 6B**, P = 0.0013) and by 40% in SJSA1 (**Figure S3A**, P < 0.001), but not in MG-63 (**Figure 6C**) or U-2 OS cells (**Figure S3B**). Interestingly, 2 mM DCA partially protected osteosarcoma cells from AA toxicity, leading to either no significant change in cell viability as in MG-63 (**Figure 6C“)** and U-2 OS **(Figure S3B)** cells, or a trend to increased cell viability, as in 143B **(“Figure 6B**, P = 0.066) and in SJSA1 **(Figure S3A**, P = 0.006) cells. However, screening 25 nM AA in the remaining osteosarcoma cell lines did not significantly impact their viability (data not shown). Prolonged (6-day) AA treatment in 17 mM glucose media in combination with 2.6 µM ONC201 and 0.25 µM ONC206 alone or together with either 2 mM DCA or 2 mM metformin mildly inhibited hFOB growth (**Figure 6D**) but completely ablated 143B cell viability (**Figure 6E“)**, and, with the exception of MG-63 in which approximately 40% reduced viability was seen (**Figure 6F**), significantly reduced viability by more than 50% in all of the other osteosarcoma tumor cell lines relative to hFOB (**Figure 6G“, Figure S3C-F).** Interestingly, adding either DCA or metformin did not lead to substantially greater cell death relative to combination ONC201 and ONC206 therapy except in Saos-2 cells (**Figure S3D**). However, the combination therapy with ONC201, AA, and DCA exhibited significant inhibition exceeding 50% reduction of viability in 6 of 7 the osteosarcoma cell lines across 4 biological replicate experiments (**Figure 6H-K“, Figure S3G-J**, P < 0.001) except for 143B, which saw only a 30% reduction in cell viability (**Figure 6I**, p < 0.001) and no toxicity in healthy hFOB cells. While RNA-Seq analysis of this combinatorial treatment was not performed, we postulate that its effectiveness may result from simultaneous targeting of unique cellular pathways. For example, metformin uniquely downregulated cholesterol biosynthesis that is crucial for cell proliferation, while ONC201 and ONC206 treatments induced apoptotic cleavage of cellular proteins (**Figure 5**). These individual treatment insights combined with experimental viability results suggest that combining different therapeutic classes in osteosarcoma may work through different cellular pathways to both prevent proliferation and induce apoptosis. This effect was demonstrated in our metastatic cell line, which was not responsive to most single-agent therapies but showed improved cytotoxicity with combination treatments (**Figure 6G,K**).

**Fig 6.**
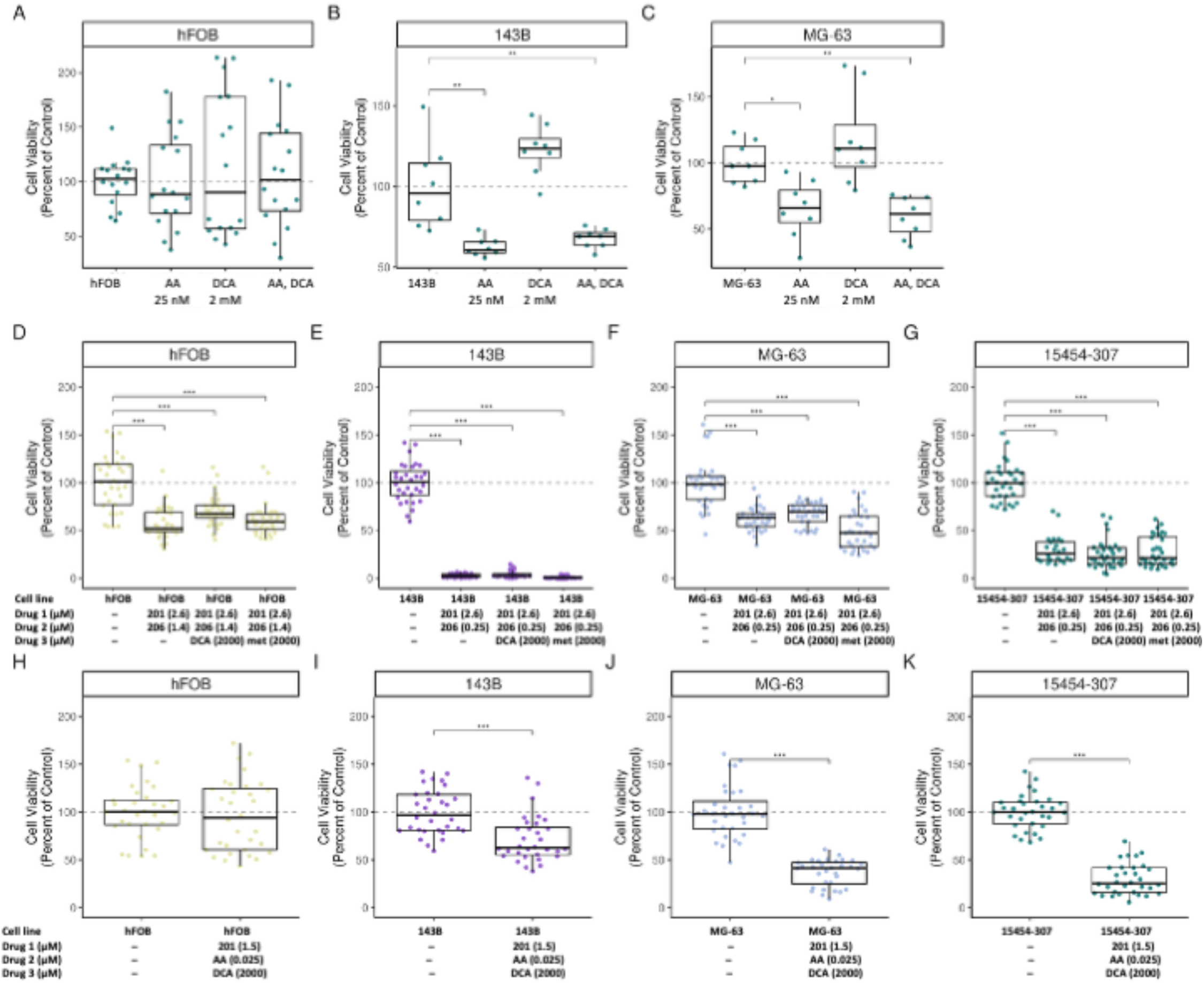
Metabolism targeting drugs alone or as combination treatments killed osteosarcoma cells but not osteoblast cell in low glucose medium for 6 days. (A-C) 2500 cells per well were seeded in 96 well plates with DMEM 10% FBS 12.5 mM glucose medium, and, after culturing overnight, the media was removed and replaced with the same medium with or without 25 nM antimycin A (AA) alone or with 2 mM dichloroacetate (DCA) alone or with a combination of the same concentrations of AA and DCA in **(A)** healthy osteoblast hFOB cells, **(B)** primary osteosarcoma 143B or **(C)** MG63 cells. **(D-G)** Cells were treated with a combination of 2.5 µM ONC201 (201) and 0.25 µM ONC206 (206), either with just the two-drug treatment or with the addition of 2 mM DCA, or 2 mM metformin (met) in 200 µL medium per well in (D) hFOB, (E) 143B, (F) MG-63 or (G) 15454-307 cells. **(H-K)** Cells were also treated with a combination of 2.5 µM ONC201, 2 mM DCA, and 2 mM met in 200 µL medium per well in (H) hFOB, (I) 143B, (J) MG-63 or (K) 15454-307 cells. After cells were cultured for 6 days, then MTT cell viability assays were performed with n = 3 biological replicate experiments. Cell viability was tested using ANOVA with a post-hoc Tukey test for all comparisons. Box plots identify the median second and third quartiles and the whiskers extend to the smallest and largest values no more than 1.5 times +/- the interquartile range. Points outside that range are considered “outlying” and are plotted individually.

### Mechanistic analysis of imipridone effects on ClpXP expression in 143B osteosarcoma cell lines

In silico analysis of public pediatric cancer variant datasets from PedCBio portal (50) suggested that ClpP was altered to a similar extent as had been previously reported in brain tumors, showing copy number amplification in a similar percentage of cases as chordomas and pheochromocytomas (**Figure 7A**). ClpX alterations were also most common in osteosarcomas over all other cancers assessed (**Figure 7B**). Based on this information and following cell viability experiments described above, further mechanistic studies were performed to investigate the connection between imipridone efficacy to reduce osteosarcoma viability and expression of the ClpXP protease complex. Specifically, hFOB and 143B cells at 90% confluence were exposed for 24 h to varying concentrations of ONC201 or ONC206. Western immunoblots of ClpP and ClpX were normalized to expression levels of CS, an enzyme within the tricarboxylic acid cycle that is frequently used as a proxy for mitochondrial content, or GAPDH that is a cytoplasmic housekeeping protein (**Figure 7C“, Figure S4 for immunoblots**). Indeed, relative to hFOB osteoblasts, 143B osteosarcoma cells had significantly increased ClpX to a greater extent than ClpP when normalized to CS (**Figure 7D-E“)**. With increasing ONC201 concentrations, a dose-dependent decrease in ClpP expression was seen **(Figure S4P,W).** In contrast, higher concentrations of ONC206 reduced ClpP expression (**Figure S4P,W**). This suggests a treatment-induced imbalance of ClpP and ClpX, which could induce the dysfunction required to trigger apoptosis and thus cell death. As transcript expression levels of ClpP and ClpX were not affected by RNA-Seq analysis (**Figure S4A)**, imipridone effects appear to be via modulating their protein levels.

**Fig 7.**
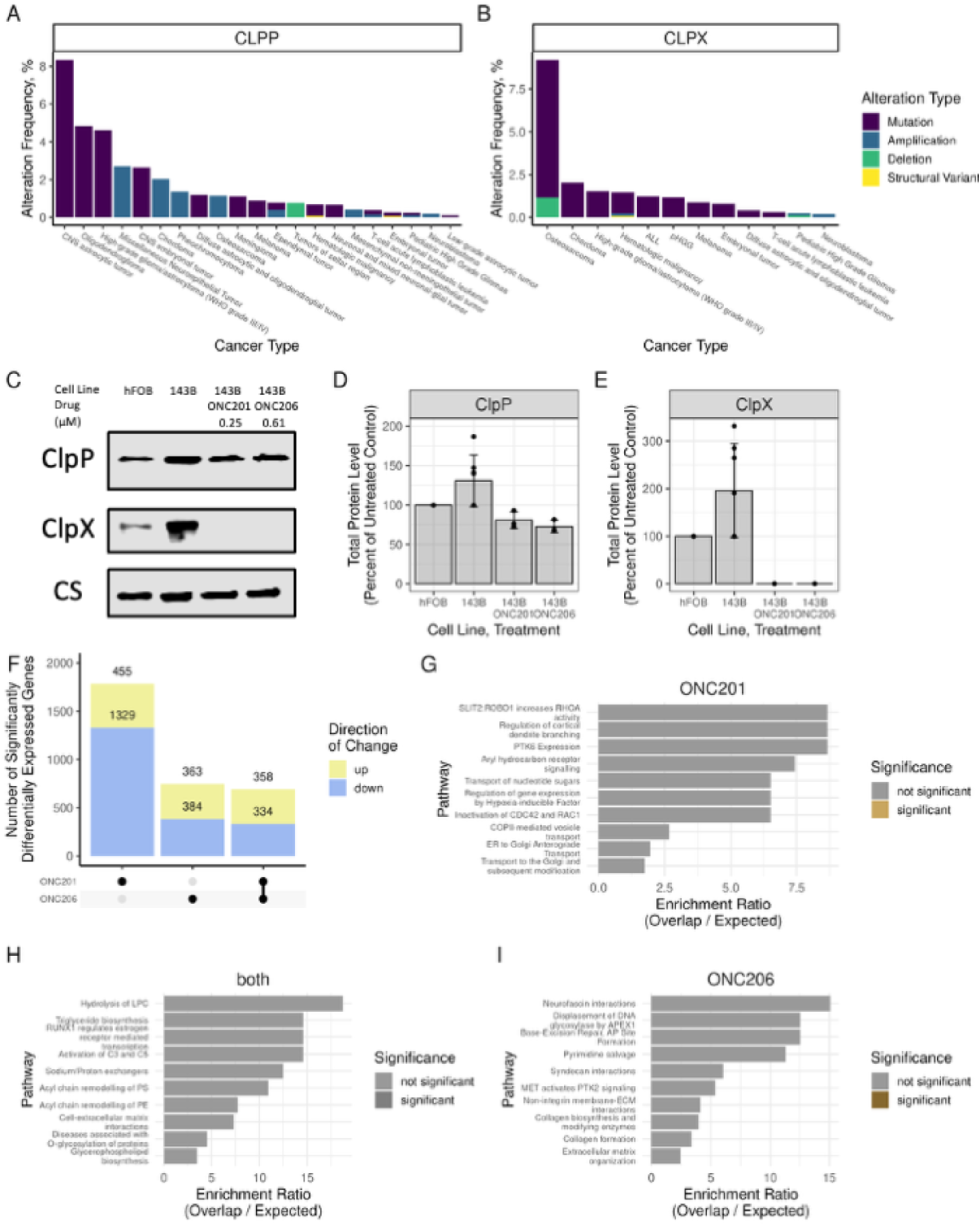
Clp alterations are frequent in osteosarcoma and ClpP imbalance induced by ONC201 or ONC206 causes distinct changes in gene expression. **(A-B)** Looking at pediatric cancers with Clp alterations, **(A)** 1.15% of osteosarcomas have ClpP alterations and **(B)** 9.20% of osteosarcomas have ClpX alterations. Cancers are given on the x-axis with the y-axis the percentage of alterations across all patients in either (A) ClpP or (B) ClpX. Color indicates the alteration type. **(C-E)** 90% confluent hFOB or 143B cells in 100 mm Petri dishes were treated with **increasing** doses of ONC201 or ONC206 indicated in 5.6 mM glucose DMEM, after 24 h, cells were collected and lysed in RIPA buffer. **(C)** Western immunoblots following SDS-PAGE (immunoblots for ClpP and ClpX are in supplemental figure S4) were imaged with anti-ClpP o**r** ClpX and anti-CS antibodies, resulting band intensities for the **untreated** and highest dose of each drug were analyzed by ImageJ, normalized to CS and relative to hFOB for ClpP **(D) and ClpX (E)**. **(F)** The differential expression results from 143B cells treated with ONC201 or ONC206 were used to evaluate the overlap in expression changes using an UpSet plot, where the bars show the number of genes changed and a single dot underneath for ONC201 and ONC206, respectively, and the line showing the number of significantly differentially expressed genes in common with both treatments. Pathway analysis, done with over-representation analysis (ORA) in WebGestaltR, on gene expression changes unique to **(G)** ONC201 alone, **(H)** ONC206 alone, **(I)** or common to both versions of the drug. Pathway analysis bar plots give the top five most up- and down-regulated pathways colored by treatment type with non-statistically significant pathways in gray **(G-I)**.

After establishing that the imipridones are working on the protein level and that the two analogues show similar effects on ClpXP protein levels in 143B cells, we went back to our RNA-Seq data to further understand the differences in effects between ONC201 and ONC206 in the same cell line. As previously mentioned, RNA-Seq transcriptome profiling of 143B cells treated with ONC201 and ONC206 transcription for common effects is evenly distributed with roughly equal numbers of genes either up or downregulated genes (**Figure 7D**). ONC206 had more specific effects vs ONC201, as it shows fewer transcriptional changes overall (**Figure 7D**). On the pathway level, no statistically significant results were seen for either the unique genes changed by ONC201 (**Figure 7E**), ONC206 (**Figure 7F**), or genes similarly regulated by both (**Figure 7G“, Table S3)**. However, going back to the full pathway analysis results for 143B cells treated with ONC201 and ONC206, both ONC versions uniquely upregulate (compared to all other drugs in the study) apoptotic cleavage of cellular proteins, telomere extension, tRNA aminoacylation and NOTCH2, suggesting that inhibition with either imipridone may induce apoptosis (**Table S3**). As the results were not conclusive on the RNA level, further mechanistic follow up work will be necessary to understand the differences in lethality between the two drugs, which may be due to off-target effects rather than their core effects on the ClpXP complex.

## DISCUSSION

Mammalian cells obtain ATP, the biochemical energy source essential for cell survival and biosynthesis, through a combination of cytosolic anaerobic glycolysis and mitochondrial OXPHOS. In contrast to healthy somatic cells, cancer cells typically obtain a greater fraction of their ATP from glycolysis relative to OXPHOS, even in the presence of sufficient oxygen (51). As anticipated, we observed that osteosarcoma 143B cells grow much faster and also have higher mitochondrial membrane potential than osteoblast (hFOB) cells of origin when grown in high relative to low glucose medium, as the latter is rate-limiting to generate ATP by glycolysis. In galactose medium where glycolysis is diminished (48),media that severely limits glycolytic capacity to generate ATP in OXPHOS compromised cells (48), almost all osteosarcoma cell lines grew more slowly than in low glucose medium, further reflecting the impaired mitochondrial OXPHOS capacity of osteosarcoma cells.

RNA-Seq transcriptome-wide profiling further demonstrated decreases upon a shift to galactose media of diverse aspects of mitochondrial metabolism, such as cholesterol and fatty acid synthesis, suggesting that decreased energy availability in the oxphos-deficient osteosarcoma cells also slows cellular production of essential compounds and slows cancer proliferation. Collectively, these data indicate that a hallmark of osteosarcoma cells is their increased dependence relative to osteoblasts on glucose to fuel aerobic glycolysis for the increased production of cellular metabolites required for generation of new biomass and to facilitate nutrient signaling. This is consistent with the Warburg classical effect in that they are highly glycolytic but further highlights their deficient mitochondrial function. Therefore, reducing glycolytic activity offers an important target for osteosarcoma therapy which may potentially be achieved either by minimizing glucose availability or using glycolytic inhibitors as complementary strategies to enhance chemotherapeutic effects and suppress tumorigenesis in osteosarcoma cells.

Given their metabolic vulnerability, extensive studies were performed to evaluate metabolic and mitochondrial modulating therapies preclinical efficacy and mechanism in osteosarcoma. As CHX inhibits cytosolic but not mitochondrial translation, we hypothesized that osteosarcoma cells that have impaired mitochondrial OXPHOS capacity would be hypersensitive to CHX treatment relative to osteoblast control cells. Indeed, prolonged treatment for 6 days with low-dose (5 µM) CHX effectively suppressed the viability of all osteosarcoma cell lines, although it also induced a lesser degree of toxicity in normal cells. RNA-Seq profiling demonstrated that osteosarcoma cells, particularly the metastatic line, treated with CHX had upregulated expression of the receptor tyrosine kinase (RTK) pathway, a known cancer proliferation pathway, which may represent a mechanism through which the surviving cells escaped apoptosis. Metformin has a controversial mechanism of action in diverse cell processes, which has variably been described as activating or inhibiting mitochondrial complex I-dependent respiration (52) and glycerol-3-phosphate dehydrogenase (53). Here, RNA-Seq studies of osteosarcoma cells suggested that metformin suppressed 143B cell growth by inhibiting the TCA cycle. However, metformin had minimal impact on other osteosarcoma cell lines’ viability, only showing efficacy in the U-2 OS cell line but not the other 6 lines tested. Thus, metformin may not be sufficient as a stand-alone therapy in all osteosarcoma cases, similarly as was identified in prior clinical trials in human osteosarcoma patients (54). A direct mitochondrial respiratory chain complex III inhibitor, AA, showed similar results as metformin when employed as a single-agent treatment, significantly decreasing cell survival in only half of the cell lines tested: 143B, MG-63, and SJSA1. Interestingly, DCA, which inhibits pyruvate dehydrogenase kinase to activate the pyruvate dehydrogenase (PDH) complex and force a metabolic switch to enhance carbon flux from glycolysis to mitochondrial respiration, had no deleterious effects as a single-agent in any osteosarcoma cell line; rather, DCA treatment yielded a trend toward increasing cancer cell viability, supporting the premise that enhancing mitochondrial function is important for cancer proliferation. As genetic heterogeneity is a well-established phenomenon in tumor cells, especially during metastatic stages (33), we postulate that the varied response to these metabolic therapies reflects in part the extensive heterogeneity known to exist between the individual osteosarcoma cell lines (55). The observed sporadic but non-universal efficacy of metabolic inhibitors suggests that targeting mitochondrial metabolism in osteosarcoma may only provide additional opportunities for the development of novel antitumor therapeutics in specific genetic backgrounds. However, their use as combinatorial treatments, including with classical anti-tumor drugs and in states of glucose restriction or glycolytic inhibition, warrants further study.

Mitochondrial ClpP is a mitochondrial serine protease that forms a complex with ClpX to regulate mitochondrial protein quality control by degrading misfolded or damaged respiratory chain proteins, thus maintaining normal metabolic function. ClpXP also regulates mitochondrial stress via induction of the UPR^mt^). ClpXP overexpression has been increasingly recognized in hematologic malignancies and solid tumors, and necessary for the viability of a subset of tumors (56). Targeting ClpP has been found to be particularly effective in glioblastoma, especially in combination treatment, for which it has recently received FDA approved (17).

Interestingly, both inhibition and hyperactivation of ClpXP have been shown to impair RC activity and induce cancer cell death (34). Remarkably, we found that not only do the imipridone class of Clp agonists kill osteosarcoma cells but they demonstrate synergistic effect when combined with metformin in 143B cells, with limited toxicity in normal osteoblasts. These findings demonstrate that targeting orthogonal mitochondrial pathways may offer a more effective anti-cancer therapeutic strategy than inhibiting single pathways. Single-agent treatment effects were found by RNA-Seq profiling to upregulate translation initiation and amino acid metabolism, which may reflect a cellular attempt to synthesize fresh copies of damaged proteins for which removal was no longer possible. Combined with pathway insights gained by RNA-Seq profiling showing that metformin reduces TCA cycle pathway activity and that imipridone treatment increases protein synthesis likely to meet energy demands and replace damaged proteins, we postulate that combination treatment with both imipridones and metformin may have greater ability to reduce cell viability in osteosarcoma cells grown in galactose by blocking both glycolytic and mitochondrial avenues for energy production as well as broader cellular opportunities to facilitate necessary metabolite and protein synthesis.

Therapeutic synergy in combinatorial treatments is highly desired, as it may permit each drug in the combination to be delivered at lower doses that minimize each drug’s (potential) side effects. While a complete characterization of the synergy of a combination therapy requires a matrix of analyses where each drug is independently varied by 2 log units across its effective dose (e.g. from 0.1 – 10 × LD_50_) (57), we cast a broader net by assaying for significant therapeutic improvements at demonstrated doses where the exhibited synergy is greatest (*i.e.,* when each drug in the combination is evaluated at ∼ half of its LD_50_). This approach allowed us to identify probable combination therapies for future careful consideration to bring forward as candidate synergistic therapies. As an example, the effect of the combination of metformin (2 mM) and ONC201 (0.35 µM) on 143B cell viability exceeded that of either agent alone. Adding metformin to ONC206 also showed a suggestion of synergy. However, the most effective synergistic therapy was a ternary therapy of metformin with both ONC201 and ONC206. This ternary therapy was also the most efficacious in MG-63 and 15454-307 cell lines.–Combining ONC 206 and ONC201 alone did not identify a significant synergistic effect in decreasing osteosarcoma viability, likely reflecting their structural symmetry, although differences clearly exist in their function given the identified differential RNA-Seq profiling responses in the osteosarcoma cells.

Synergy among metabolic modifying therapies may also have limits, as no synergy with imipridones was observed when combined with AA or DCA in any of the four cell lines tested. However, DCA and ONC201 together when combined with either ONC206 or AA, yielded relatively enhanced cytotoxicity compared to single-agents in multiple osteosarcoma cell lines, with good therapeutic index based on minimal toxicity to normal osteoblast cells.

Overall, our study highlights the significant influence of the tumor nutrient environment on cancer cell behavior, demonstrating distinct effects in osteosarcoma from those driven by the osteoblast somatic cell of origin. We further identified the therapeutic potential in osteosarcoma of metabolic modulating chemotherapeutic agents (CHX that inhibits cytosolic translation and imipridones that modulate ClpXP) that have not been previously explored in osteosarcoma, showing great variability in their effectiveness as single agents but synergistic effects when combined with strategies that modulate mitochondrial activity either through cellular nutrient manipulation (glucose restriction and galactose-only exposure) or small molecule modulating therapies (metformin, AA). Conversely, DCA single-agent therapy to increase mitochondrial metabolism via enhancing PDH activity had no toxic effect but instead appeared to enhance osteosarcoma proliferative capacity. These findings not only support the repurposing for osteosarcoma that has no curative standard and high metastatic rate of existing drugs such as imipridones, but also pave the way for future investigations aimed at exploiting the demonstrated impairment in mitochondrial OXPHOS capacity of osteosarcoma cells as combination therapy approaches with standard chemotherapies. The merit of this proposed combinatorial metabolic modulation plus chemotherapy therapeutic approach is underscored by the strongly favorable therapeutic index of this class of therapies, enabling the potential dose reduction of chemotherapies with toxic profiles. Collectively, this work offers a compelling preclinical foundation for novel therapeutic metabolic approaches that may be pursued in future rigorous clinical trials to improve survival and outcomes for osteosarcoma patients.

## AUTHOR CONTRIBUTIONS

MJF and AR conceived of and designed the study and obtained project funding. MP, with the assistance of SD, performed cell line culture, MTT viability assay, CX5 mitochondrial physiology assay, immunoblot, RNA-Seq sample preparation, and data analyses, and prepared figures. KK analyzed all RNA-Seq data and plotted all figures. VEA contributed to the figures, discussion, and provided editorial assistance. MP, SD, KK, and MJF wrote the paper, with methods sections prepared by each author who performed the respective experiments. All authors approved of the final version.

